# AnoniMME: Bringing Anonymity to the Matchmaker Exchange Platform for Rare Disease Gene Discovery

**DOI:** 10.1101/262295

**Authors:** Bristena Oprisanu, Emiliano De Cristofaro

**Affiliations:** University College London

## Abstract

**Motivation:** Advances in genome sequencing and genomics research are bringing us closer to a new era of personalized medicine, where healthcare can be tailored to the individual’s genetic makeup, and to more effective diagnosis and treatment of rare genetic diseases. Much of this progress depends on collaborations and access to data, thus, a number of initiatives have been introduced to support seamless data sharing. Among these, the Global Alliance for Genomics and Health has developed and operates a platform, called Matchmaker Exchange, which allows researchers to perform queries for rare genetic disease discovery over multiple federated databases. Queries include gene variations which are linked to rare diseases, and the ability to find other researchers that have seen or have interest in those variations is extremely valuable. Nonetheless, in some cases, researchers may be reluctant to use the platform since the queries they make (thus, what they are working on) are revealed to other researchers, and this creates concerns with respect to privacy and competitive advantage.

**Contributions:** In this paper, we present AnoniMME, a framework geared to enable anonymous queries within the Matchmaker Exchange platform. The framework, building on a cryptographic primitive called Reverse Private Information Retrieval, let researchers anonymously query the federated platform, in a multi-server setting—specifically, they write their query, along with a public encryption key, anonymously in a public database. Responses are also supported, so that other researchers can respond to queries by providing their encrypted contact details.

**Availability and Implementation:** https://github.com/bristena-op/AnoniMME.

## 1 Introduction

Advances in genome sequencing and genomics are enabling tremendous progress in medicine and healthcare, paving the way to making the prevention, diagnosis, and treatment of diseases tailored to the individual’s specific genetic makeup, thus becoming cheaper and more effective. Researchers are also gaining a better understanding, and developing more successful treatments of rare genetic diseases. However, even though sequencing costs have plummeted from billions to thousands of dollars over the past 15 years (see https://www.genome.gov/sequencingcosts/), it is still hard for researchers to gain access to genomic data, especially those pertaining to rare conditions.

Therefore, seamless progress in genomics research hinges on the ability to collaborate and share data among different institutions. Indeed, funding agencies often require that data sharing is considered in grant applications, and a number of initiatives have been announced to gather and share genomic data. For instance, the All Of Us Research Program (formerly known as the Precision Medicine initiative) was launched in the US in 2015, aiming to collect health and genetic data from one million citizens. Similar projects exist elsewhere, e.g., in the UK, Genomics England is sequencing the genomes of 100,000 patients, focusing on rare diseases and cancer. There are also initiatives specifically targeting data sharing, such as the NIH’s Genomic Data Commons (GDC), which provides the cancer research community with a unified data repository across cancer genomic studies (see https://gdc.cancer.gov/).

Aiming to foster collaborations, the Global Alliance for Genomics and Health (GA4GH) [https://www.ga4gh.org] was established, with core funding from NIH, Wellcome, and Canada’s CanShare, with the explicit goal of making data sharing between institutes simple and effective. The GA4GH has developed several platforms, e.g., the Beacon Project [4], allowing researchers to search if a certain allele exists in a database of genomic data, as well as the Matchmaker Exchange (MME) [8], which facilitates rare disease discovery.

In this paper, we focus on the latter; The MME platform connects multiple distributed databases through an API and allows researchers to query for genetic variants in other databases in the network. That is, MME acts as a portal supporting simultaneous querying over multiple databases that are members of the exchange. More specifically, MME allows a researcher to query a specific gene, e.g., “AP3B2” (a gene where rare mutations have been linked to early-onset epileptic encephalopathy). If a match is found, the researcher is notified of all matches within all databases in the MME, and can get in touch with the user that submitted the case on which a match is generated. Note that, querying a gene really implies querying a known rare variation of that gene.

However, researchers might be reluctant to use the platform since the queries they make are revealed to other researchers, and this exposes what they are working on and what kinds of patients they might have, ultimately resulting in loss of privacy and competitive advantage. Indeed, MME currently requires researchers to submit a registration application to be given access to the platform, with the goal of preventing misuse of the system, thus, queries made on this platform are not anonymous and are revealed to all other researchers with an interest in the same gene.

### Problem Statement

This motivates the need to support *anonymous querying* on MME, so that a researcher’s interest a gene is not broadcast, but only communicated to relevant contacts, i.e., researchers with same interests or willing to collaborat-e. To this end, we present AnoniMME, a framework letting researchers anonymously query a gene within the MME, without violating any of MME’s current functionalities and requirements. We build AnoniMME using a cryptographic primitive called Reverse Private Information Retrieval, using a model similar to that presented by the anonymous messaging system Riposte [3], while creating queries and implementing the same functionalities as in MME. In other words, researchers can perform anonymous queries to the federated platform, in a multi-server setting, by writing their query, along with a public encryption key, anonymously, in a public database. AnoniMME also supports responses, so that other researchers can respond to queries by providing their encrypted contact details.

### Solution Intuition

We build queries in regular epochs, where the length of each epoch is based on the number of write requests. In order to anonymously write to the database, the user selects a random row of the the database, and splits the query, containing the gene and her public key, into shares, one for each server (which we denote as *node* servers). This way, the node servers cannot learn anything about the write request, if at least one of the them is honest. Then, a *master* server can gather queries that have been collected during an epoch from the node servers and collate them together to recover and publish the actual queries. The MME matching system can then be used in order to generate matches for the queries, in the usual manner, and contact details of other researchers/clinicians can be exchanged, encrypted using the public key, and published in the same row as the queried gene, in an adjacent column.

To demonstrate the practicality of AnoniMME, we implement and evaluate our prototype experimentally. We do so in two different settings, one involving two node servers and a master server, and another involving six node servers (and a master server). In both settings, the nodes collect write requests during an epoch, and then forward them to the master server which collates them and publishes the final database.

### Contributions

In summary, our paper makes several contributions:

1. We present AnoniMME, a framework enabling anonymous queries within the Matchmaker Exchange (MME), without breaking any of its current security and functionality requirements.
2. We build AnoniMME from Reverse PIR [3], using an information-theoretic approach, extending queries to support public key encryption of contact details, and adding a response phase so that users can also anonymously reply to queries.
3. We show, experimentally, that AnoniMME is efficient and scalable, and can bring anonymity to MME with low overhead. Therefore, we are confident that it can be deployed in the wild and further encouraging researchers to share genomic data.

### Paper Organization

The rest of the paper is organized as follows. In the next section, we introduce our approach; specifically, after reviewing the Matchmaker Exchange (MME) platform, we define entities, operations and threat model of our system, and present a first attempt at designing a anonymous-query mechanism for MME. In Section 3, we then describe the methods used for collision handling and collision recovery, present the *n*-server protocol, and evaluate the performance of the proposed protocol on the client side. Next, in Section 4, we discuss the results from our experimental evaluation and place our protocol in the context of related work. Finally, the paper concludes in Section 5.

## 2 Approach

### 2.1 Matchmaker Exchange

As mentioned, the Global Alliance for Genomics and Health (GA4GH) was established, in 2013, aiming to support simple mechanisms for sharing data between institutes. The GA4GH has developed and deployed various systems, including the Matchmaker Exchange (MME) [8], which facilitates rare disease gene discovery and constitutes the main focus of our work. MME is a federated platform that facilitates the identification of cases with similar phenotypic and genotypic profiles through a standardized Application Programming Interface (API). Essentially, it enables searches in multiple databases, without having to query all of them separately or deposit data in each of them. As of March 2018, it involves seven organizations with full member status (AGHA Patient Archive, DECIPHER, GeneMatcher, Matchbox, Monarch Initiative, My-Gene2, and PhenomeCentral), and eight additional participant organizations.

The Matchmaker Exchange Application Programming Interface (MME API) [1] fully specifies the data format and the protocol for querying databases to identify individuals with similar phenotypic profiles and genetic variations. To ensure the accuracy of the patient comparison, similar phenotypes are determined by matching identical or ontologically similar with the Human Phenotype Ontology (HPO). The MME API also specifies the format of both the query, which is sent to participating databases (called “matchmaking service”) and the response, which contains information about matching individuals in the remote database. It is implemented under a query-by-example methodology: a user can query a specific gene, e.g., “AP3B2,” and she will be notified of all matches within all databases in the MME. Note that querying a gene really implies querying a known rare variation of that gene. If a match is found, the user receives a Case ID for the match, information about the user that submitted the case on which a match is generated, such as name, institution and email address, as well as the corresponding candidate gene or phenotype. In order to query the platform, users must be registered with one of the member databases, and have a clinician/researcher account. Some of the member databases allow for patient/family registrations as well, however, the submissions made by these type of users are excluded from matching via MME, due to the current MME rules.

The query protocol is illustrated in Figure 1. A user, Bob, sends the metadata (i.e., Case ID, submitter information) as well as the patient data (gene and/or phenotype) to Database B. Another user, Alice, submits a similar case to Database A; Database A then sends an MME API match request to Database B, which performs the match and returns a list of scored patients, along with relevant metadata, to Database A. After receiving the match results, Database A informs Alice, providing contact information for Bob. The result of querying MME yields a list of matches, where each match has a *patient* object, i.e., the information on the matched patient, consisting of the same information as described in the query, and a *score* object. The scoring of the patients is done according to how well the results patient matches the query patient, i.e., it is a numerical value in the range [0, 1], where 0.0 is a poor match and 1.0 a perfect match.

**Figure 1:**
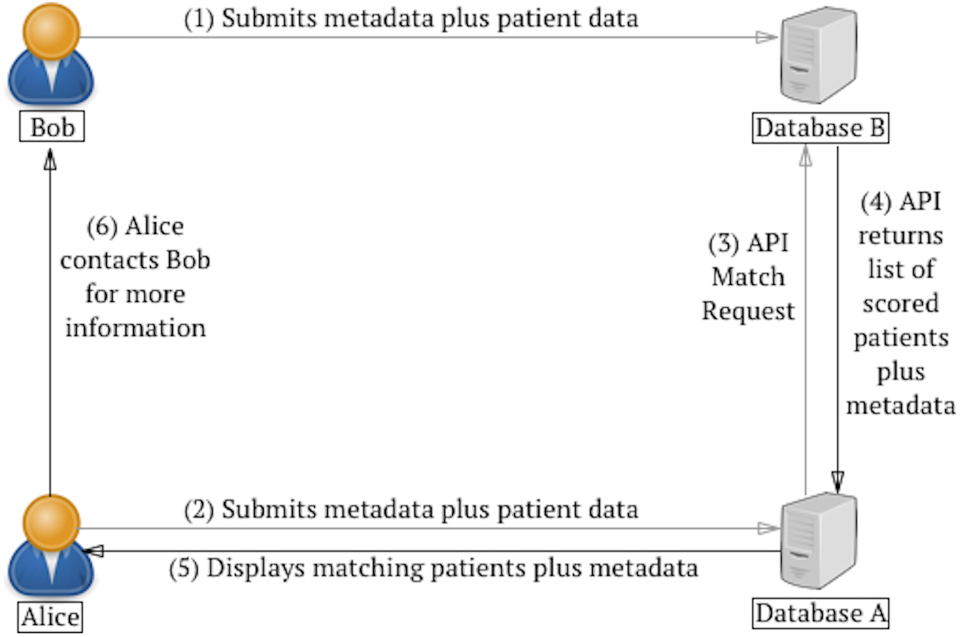
Visual representation of a MME query sequence.

### 2.2 Entities and Operations

As discussed in Section 1, this paper presents AnoniMME, a framework geared to enable anonymous queries within the Matchmaker Exchange platform, i.e., anonymously querying the federated platform to find patients with similar gene mutations or phenotypes. It involves the following entities:

#### Querying Users

researchers/clinicians who query the system to find other users that have patients with a rare mutation or an interest in the same gene. As discussed later, they generate a write request specifying the row at which their query, i.e., the gene of interest and their public key, will be processed.

#### Responding Users

researchers/clinicians replying to an existing query. They use the public key of a querying user to encrypt their contact details and generate a write request for the same row as the gene of interest including their (encrypted) contact details.

#### Nodes

the servers collecting write requests from the users. These are aggregated until the end of an epoch, based on the maximum number of write requests. Each node server can be run by one of the current MME members.

#### Master Server

a server that gathers the databases from each node at the end of an epoch, and publishes the database with all the write requests revealed. The master server role can also be assigned to one of the existing MME members, and can be reassigned to another member at the end of each epoch.

Overall, AnoniMME implements the following operations:

#### Query Write Request

On input row *i*, query gene *X*, and public key *PK*, a querying user generates *n* write requests, one for each node. Each write request is generated by encoding the gene and the public key into *n* vectors, so that all of them combined will write the gene/public key at index *i*.

#### Query Response Request

On input row *i*, encrypted contact details *c*, a responding user generates *n* write requests, one for each node. Write requests are generated, once again, by encoding the encrypted contact details into *n* vectors.

#### Database Collation

On input *n* databases, the master server collates them into one final database, and publishes it.

### 2.3 Security Model

AnoniMME aims to guarantee the following three security goals:

*1. Correctness*. When all nodes execute the protocols correctly and send data to the master server at the end of an epoch, the resulting database contains all the write requests processed as if the requests were directly applied to the final database.

*2. Anonymous Write*. The probability that an adversary guesses at which particular row a user has written is only negligibly better than random guessing.

*3. Disruption Resistance*. An adversary controlling *n* users can make at most *n* write requests (i.e., there is a limit to the number of write requests each user can make during an epoch).

#### Threat Model

We assume that the users of the system are untrusted, and may collude with the nodes, the master server, or other users in order to violate the security properties of the system. Both the master server and the nodes are trusted for availability and to follow the protocol correctly, under the assumption that at least one of the nodes is honest (i.e., does not collude with other nodes). We do not consider external adversaries, since their actions can be mitigated via standard network security techniques (i.e., using a secure and authenticated communication channel). Finally, note that the security model of AnoniMME mirrors that of Riposte [3].

### 2.4 A First Attempt

We now present a first attempt at instantiating AnoniMME, and discuss its limitations, which we address in the actual construction of AnoniMME presented in Section 3.2.

#### Intuition

We start by attempting to build from a simple extension of Reverse Private Information Retrieval (Reverse PIR) [3]. More specifically, we implement the query phase using the same mechanism of Riposte, i.e., we let users anonymously submit the gene of interest, along with their public key, with a “write request.” We then add a response phase, allowing users with an interest in the same gene to respond— specifically, by encrypting their contact information using the public key contained in the query, and adding it to another write request.

In the following, we present a construction assuming the presence of 2 servers (*S*_1_ and *S*_2_) and a database with *l* rows.

#### Query phase

Assume user A wants to anonymously query gene *X*_*A*_. She builds a write request, consisting of (*X*_*A*_, *PK*_*A*_), where *PK*_*A*_ is her public key, and writes this at row i in the database. More specifically, she picks 2*l* random numbers, *r*_1_,*r*_2_,…, *r*_*l*_ and *s*_1_,*s*_2_,…, *s*_*l*_, where *l* is the size of the database. The query write request vectors are constructed as follows:

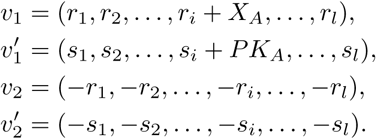

Note that *v*_1_ + *v*_2_ = *X*_*A*_ ⋅ *e*_*i*_, and 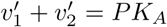, where *e*_*i*_ denotes the unit vector with 0’s at all positions except at position *i*, where it is equal to 1, and thus the construction is correct. Then, A sends 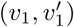 to *S*_1_, and 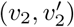 to *S*_2_.

Write requests are collected until the end of an epoch, when the servers combine their local states and publish the database with the queries. As long as the two servers do not collude, none of them can reconstruct what any given user has written, i.e., none of the servers can recover the gene or public key of the user sent in the write request. Also, in order to achieve disruption resistance, one can limit the number of queries to one per user for each phase of the epoch.

#### Response phase

After the database with the queries is published, the response phase begins. Here we can rely on MME’s algorithm to generate matches on existing MME data, and simply extend it to encrypt the contact details of the relevant users with an interest in the same gene. This would be inline with the current privacy policy of the MME, as contact details of researchers with an interest in the same gene are already shared.

Users can also be given an option to voluntarily provide their contact details as follows. If user B notices that another researcher (user A) has an interest in the same gene X, say at row i of the database, she gets A’s public key *PK*_*A*_, and encrypt her contact information (*C*_*B*_) under *PK*_*A*_ and generates a write request as a share of *Enc*_*PKA*_(*C*_*B*_), in a similar manner to the first epoch. More specifically, she chooses random 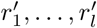 and forms the following vectors:

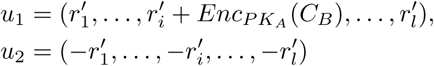

User B then sends *u*_1_ to server *S*_1_ and *u*_2_ to *S*_2_. At the end of this epoch, the results are being published in a column adjacent to the queried gene and the public encryption key. The querying users can use the database to find the row of interest (in this case *i*), decrypt the contact details, and get in touch with the responding users.

#### Correctness and Security

It is straightforward to see that the construction is correct, since, if all nodes execute the protocols correctly the result of combining all their local database states at the end of an epoch by the master server will result in revealing all the write requests processed. An adversary’s advantage of guessing at which a certain user has written in the final database is the same as random guessing, hence, the construction guarantees anonymous writes. Disruption resistance can be also achieved in a straightforward manner since MME requires users to register on one of the databases, so they can allow maximum one write request per registered user per epoch.

#### Limitations

Alas, this construction has the following limitations:

1. *Collisions*: They might occur for writes generated by honest users, which all want to write at the same row;
2. *Maliciously-formed write requests*: A malicious user can easily send a malformed request to the servers, making all the data within the database non recoverable.

## 3 Methods

In this section we provide methods for collision handling for our first attempt and use it to provide a description of the n-server protocol. We also evaluate the proposed method in terms of time and bandwidth required in order to asses the feasibility of the proposed construction.

### 3.1 Handling Collisions

As discussed previously, collisions might occur whenever multiple users want to write at the same row. Aiming to address them, we set the database size to be large enough to accommodate write requests at a 95% non-collision rate. In other words, 5% of the queries will likely fail due to collisions and will need to be re-submitted.

#### 3.1.1 Minimizing collisions

Our intuition is to follow a “balls and bins” approach, i.e., if we throw m balls uniformly and randomly into the *l* bins, we can estimate how many bins will contain exactly one ball. In our model, we can associate write requests to the *m* balls and the rows of the database to the *l* bins. Let *B*_*ij*_ be the event that ball i falls into bin *j*: for all *i* and *j*, we have 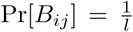. Then, let 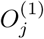 be the event that exactly one ball falls in bin *j*. We have that:

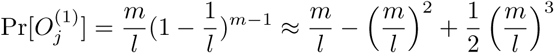

using the binomial theorem and ignoring low order terms. Then, 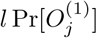 is the expected number of bins with exactly one ball, i.e., the expected number of messages successfully received. Dividing by *m*, we get the expected success rate as

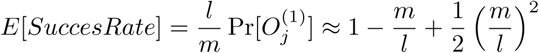

Thus, for a 95% expected success rate, we need *l* ≈ 19.5*m*.

In AnoniMME, in order to set the size of the database, we need to estimate the expected number of write requests for each epoch. Looking at the three MME members which show statistics on the number of users, we find that GeneMatcher has 4, 066 registered users, MyGene2 345 registered families, and Decipher 247 registered projects (users have to be part of a project in order to join Decipher) as of November 2017. This yields an average of approximately 1, 550 users per database. Assuming that this is representative of the number of users for all MME databases, we can approximate the total number of users to be in the order 10, 000. We also need to estimate how many users make queries in each epoch: assuming 5% of users do so at each epoch, each epoch can run for 500 queries, yielding a database of size *l* ≈ 10, 000. Further, note that we design AnoniMME’s write request so that the row number at which we write is determined at random, given the number of write requests in the epoch as well as the database size, in order to avoid biases in choosing rows. This method, however, does not provide any way to recover in the case where a collision occurs, in that case the queries are irrecoverable, and the users would need to resubmit their queries in a future epoch.

#### 3.1.2 Recovering from collisions

We also use a simple technique for recovering from collisions if/when these occur. Assume *α* messages have been written at row *i*, i.e., we have *a* = *m*_1_ + *m*_2_ + … + *m*_*α*_. Inspired by [3], we can modify the way in which the queries are built to recover each of the individual message *m*_*j*_, for 1 ≤ *j* ≤ *α*; specifically, we can use a system of *α* equations, which allows us to solve for each of the colliding messages. Without loss of generality, we consider the case *α* = 2 and explain how to recover from collisions occurring for the gene name, but similar methods can be used for *α* > 2 and to recover public key and/or encrypted contact details. When a collision occurs at row *i*, we have an entry *a* = *X*_*A*_ + *X*_*B*_, where *X*_*A*_ is the gene sent by user *A*, and *X*_*B*_ is the gene sent by user *B*. If, rather than just sending the queried gene *X,* users send (*X*, *X*^2^), we can recover *X*_*A*_ and *X*_*B*_ by solving a system of two equations with two variables.

In this case, we also compute the size of the database needed for an expected success rate as follows:

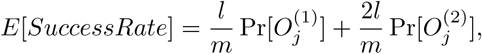

where 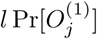 is the expected number of rows with exactly one write request applied to them, computed as before, and 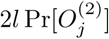 is the expected number of rows with exactly two write requests applied to them. Computing 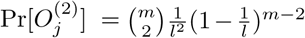, we obtain the value of the expected success rate as:

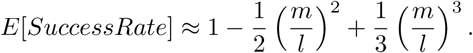

In this case, for an epoch of *m* write requests, with a 95% expected success rate, we need a database with *l*′ ≈ 2.7 *m* cells (two columns and 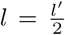 rows). This implies that with 500 write requests per epoch, the database needs *l*′ ≈ 2.7-500 = 1,350 cells for each vector.

We now generalize for any value of *α*. Users submit *X*, *X*^2^,…, *X*^*α*^ for any gene *X* to be queried. This allows us to recover from an *α*-way collision as, in that case we obtain a system of *α* equations with *α* variables. The expected success rate is:

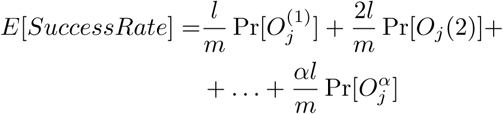

where 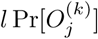 is the expected number of rows with exactly *k* write requests applied to them. Each 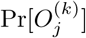 is computed as 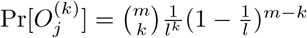. Hence, we obtain:

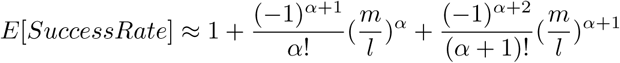

We solve this equation for *l*, given the expected success rate *E*[*SuccessRate*], the collision recovery factor *α* and *m* the number of write requests to be written in a certain epoch. If this method is used throughout both epochs, colliding requests from the query phase will have to be recovered before the response phase can begin.

Due to the nature of our query/response model, we can expect collisions to occur more often in the response phase. Hence, we will build the system using different collision recovery factors *α*_*q*_ for the query phase and *α*_*r*_ for the response phase, with *α*_*r*_ ≥ *α*_*q*_.

### 3.2 N-server Construction

We now present the generalized model for the case with *n* servers and a database with *l* rows. We use collision parameters *α*_*q*_ and *α*_*r*_ for the query and response phase, respectively. The various steps of the construction are illustrated in Figure 2.

**Figure 2:**
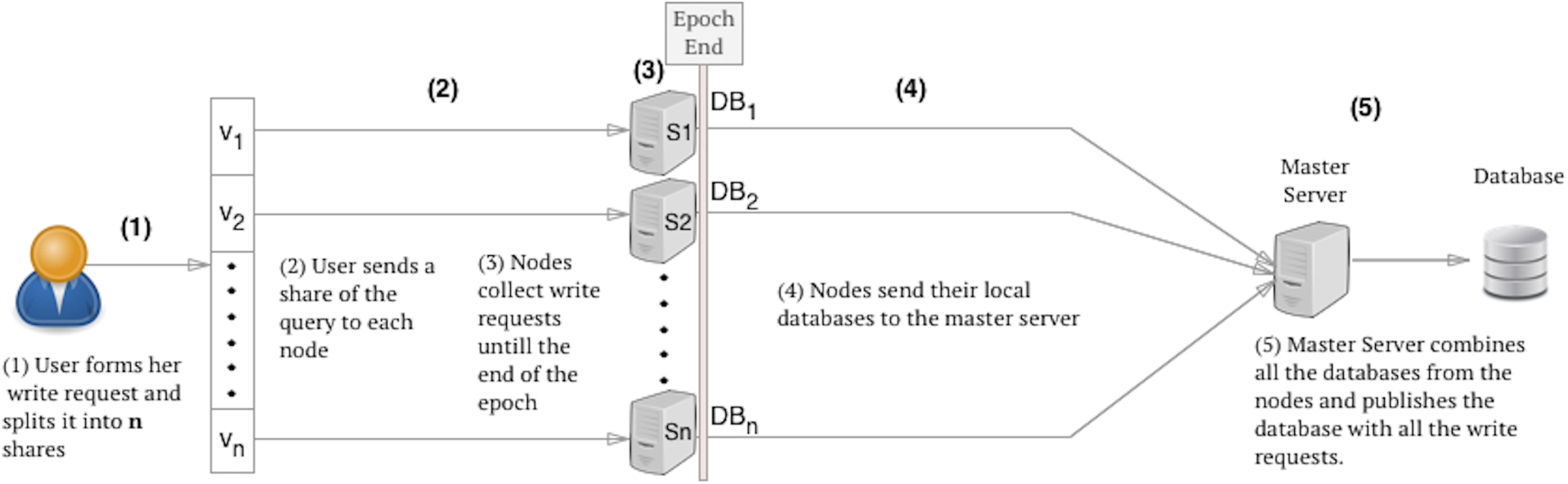
*n*-server write request processing. At the end of the epoch the Master Server publishes the database with all the write requests and the nodes will be reset to hold an empty database.

#### Query phase

Assume user A wants to query gene *X*_*A*_, but does not want to reveal that she is the person querying it. As in the construction presented in Section 2.4, A builds her write request, consisting of (*X*_*A*_, *PK*_*A*_), where *PK*_*A*_ is her public key, aiming to write at row *i* in the database. She picks random numbers 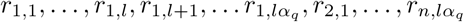 and 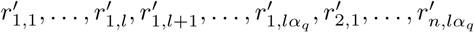, where *l* is the size of the database, *n* the number of nodes the write request will be sent to, and *α*_*q*_ the number of allowed collisions. The query write request vectors are then constructed as follows:

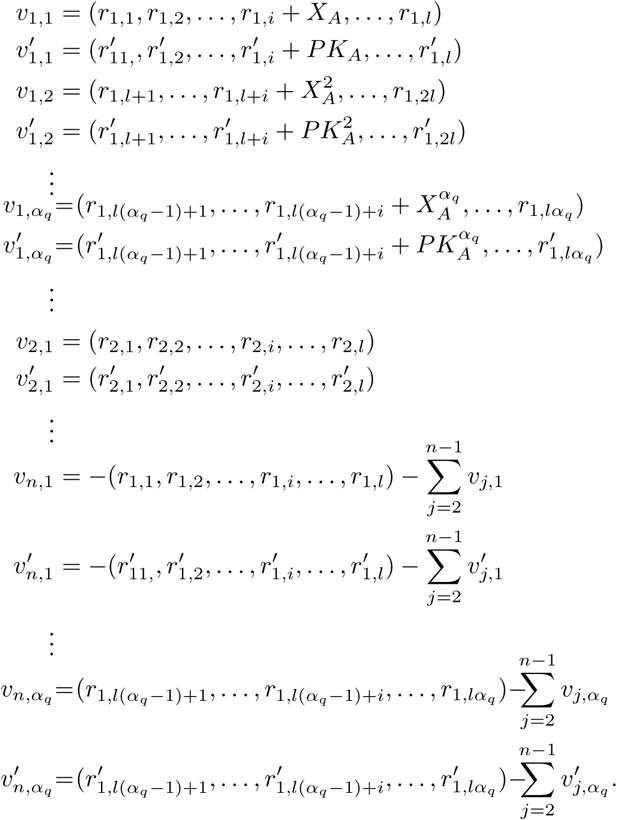

The querying user A ends 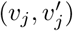 to server *j* for each *j*, 1 ≤ *j* ≤ *n*, where *V*_*j*_ = (*V*_*j*_, 1, …, *v*_*j,α*_*q*__) and 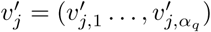. We also consider the special case of *α*_*q*_ = 1, when there is no recovery for collisions, but, instead, we adjust the database size according to the minimizing collisions case. The servers collect write requests until the end of the epoch and then send their local databases to the master server, which will combine them to reveal the database.

#### Response Phase

As the database with the queries is published, the response phase begins. As discussed in Section 2.4, we can rely on MME’s algorithm to generate matches on existing data from the platform, encrypt the contact details of the relevant users with an interest in the same gene, and extend it to allow for voluntary responses. More specifically, user B can add their contact details *C*_*B*_ by sending a write request as a share of *c* = *Enc*_*PK A*_ (*C*_*B*_), in a similar manner to the first epoch. That is, first, she picks random *s*_1,1_, …, *s*_1,l_, *s*_1,*l* + 1_, …, *s*_1,*lαr*, *s*_2,1__, …, *s*_*n,lα*_*r*__ and forms the following vectors:

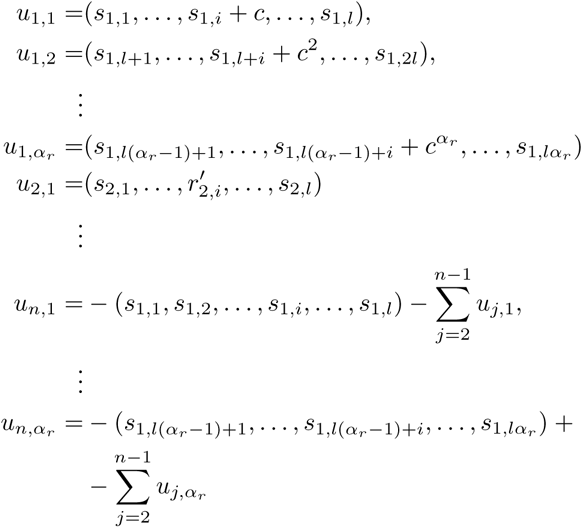

User B then sends *u*_*j*_ = (*u*_*j*,_1__,… *u*_*j*,*α*_*q*__) to server *S*_*j*_. At the end of this epoch, the results are being published in a column adjacent to the queried gene and the public encryption key. In case of collisions, the individual ciphertexts can be recovered up to *α*_*r*_ collisions. Finally, the querying users can use the database to find the row of interest (in this case *i*) and decrypt the contact details received and contact the person.

### 3.3 Experimental Evaluation

We now present an experimental evaluation of AnoniMME, aiming to demonstrate its practicality for real-world deployment.

We have implemented the *n*-server construction (Section 3.2) using Python 3.6 and evaluated our prototype on a Macbook Pro running MacOS Sierra 10.12.6 and equipped with a 2.7GHz Intel i5 processor, and 16GB of RAM. Experiments are performed in two different settings, with two and six node servers, respectively, and always averaged over 1,000 executions. We also use three different epoch sizes, namely, 100, 500, and 1,000 write requests per epoch during the query phase. For the response phase, we keep the database size fixed from the query phase. Overall, we evaluate running times needed to generate the write requests and the bandwidth overhead supporting the recovery of 2, 5, and 10 colliding messages, all on the client side (i.e. one request per epoch).

The servers run Flask with RESTful interface, so we use HTTP requests to send the messages, and the payload is built in JSON, therefore, we measure, in bytes, the size of the JSON payload (plus HTTP headers) to estimate the total bandwidth required for sending write requests.

On the client side, the cryptographic layer includes generating public/private keys (done only once) and building the vectors to be sent to the *n* servers as part of the write request, which incurs O(*n*) complexity. Gene name and contact details are assumed to be no longer than 64 characters, while random numbers used for vector generation during query phase are up to 1,024 bits long, for *α*_*q*_ ∈ {1, 2} and *α*_*r*_ = 2. For the response phase, the length of the random values varies according to the collision recovery factor *α*_r_. For *α*_*r*_ = 5, their length is 2,560 bits, while for *α*_*r*_ = 10 it is 5,120.

Finally, note that plausible gene queries are generated using the set of gene symbols (e.g, “BRCA2”) from http://gfuncpathdb.ucdenver.edu/iddrc/iddrc/data/officialGeneSymbol.html.

#### 3.3.1 Two Node Servers

We start with the setting involving two node servers and a master server, considering epochs of size 100, 500, and 1,000. As mentioned above, we evaluate bandwidth overhead and running times required for query and response write requests.

The database size required for each of the three test cases is calculated according to the method presented in Section 3.1 for minimizing collisions, thus, *l* = 19.5*m*, where *l* denotes the number of rows required and m is the number of write requests for the epoch. It follows that the l amounts to 2,000, 10,000, and 20,000 rows for m equal to 100, 500, and 1,000, respectively.

Running times for both the query write and the response (considering *α*_*r*_ ∈ {2, 5,10}) are shown in Figure 3. Overall, we find that, during the query phase, with a database size of 2,000 rows, it takes approximately 0.014s to generate vectors in our testbed. Running times scale linearly, i.e., it takes 0.062s with 10,000 rows and 0.126s with 20,000 rows. The bandwidth overhead, shown in Figure 4, ranges from 2.5MB for the smallest database size to 25MB for the largest case considered in our test cases, which can be considered an acceptable amount of traffic expected from the client side.

**Figure 3:**
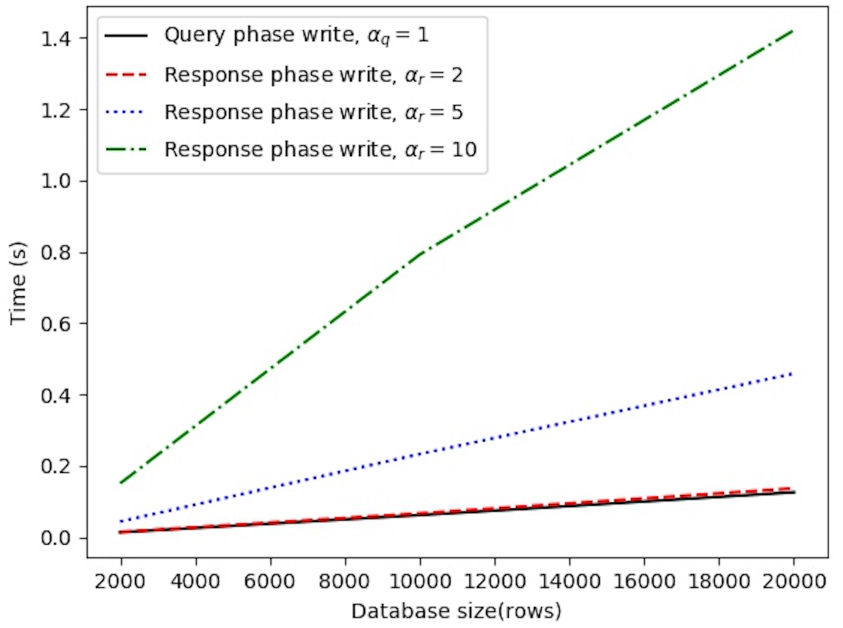
Two nodes running times for query write request, response write request with recovery from 2 collisions, response write request with recovery from 5 collisions, response write request with recovery from 10 collisions.

**Figure 4:**
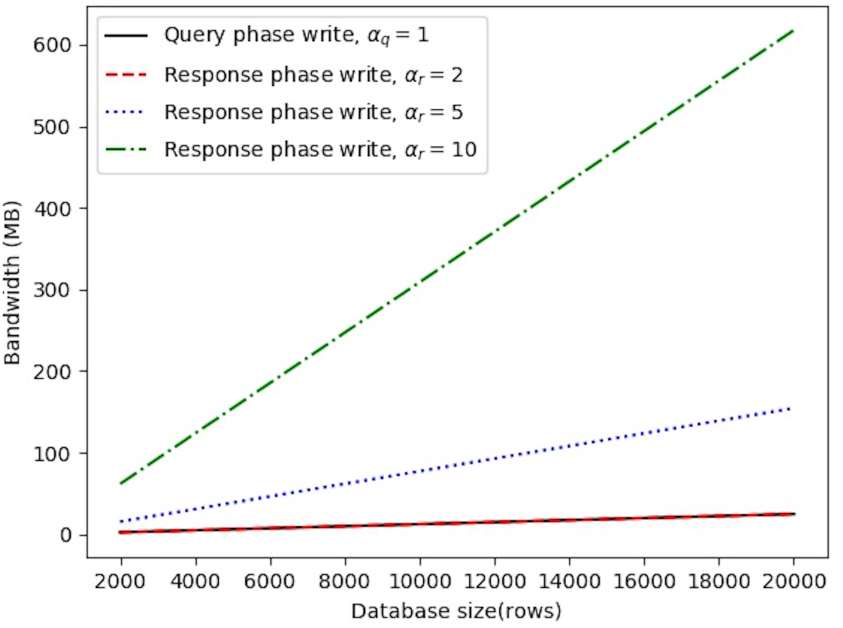
Two nodes bandwidth averages for query write request, response write request with recovery from 2 collisions, response write request with recovery from 5 collisions, response write request with recovery from 10 collisions.

For the response phase, we find that, when *α*_*r*_ = 2, the results are similar to the query phase since responding users need to generate two vectors in order to allow collision recovery, same as for the querying user. When *α*_*r*_ equals 5 or 10, we notice an increase in both running times and bandwidth. Nonetheless, computational complexity is still acceptable, since, even with the largest database size, write request generation takes less than 0.5s for *α*_*r*_ = 5 and less than 1.5s for *α*_*r*_ = 10. Communication overhead, on the other hand, increases to 160MB and 617MB, respectively, with the largest database size.

However, one can adjust the collision minimization parameter so that 10-way collision recovery is not needed.

#### 3.3.2 Six Node Servers

We also experiment with an instantiation of AnoniMME using six node servers, thus mirroring the current MME setting, which involves seven members. Once again, we consider three settings (100, 500, and 1,000 write requests per epoch), and obtain the resulting database size based on the recovery from collisions method discussed in Section 3.1. We support recovery from two colliding messages for the query phase, *α*_*q*_ = 2. Therefore, the number of rows required is 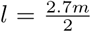, where *m* is the number of write requests for the epoch, thus, *l* equals 135, 675, and 1,350 for *m* = 100, 500, and 1,000, respectively. As per the response phase, we run tests with different values *α*_*r*_ ∈ {2, 5,10}, considering the database size fixed as for the query phase.

Once again, we estimate running times (see Figure 5) and the bandwidth overhead (see Figure 6). Even though this requires more vectors to be generated by the users compared to the two-node setting (cf. Section 3.3.1), we observe a considerable decrease in both running times and bandwidth overhead for the same epoch sizes due to the decreased number of rows in the database. Specifically, computational complexity is again linear over all test cases, but the write request generation taking less than half the time. There is also a big improvement in terms of communication complexity: even in the most bandwidth-heavy case (i.e., *α*_*r*_ = 10), with 1,000 write requests per epoch, we observe a five-fold improvement, with bandwidth decreasing from 617MB to 125MB.

**Figure 5:**
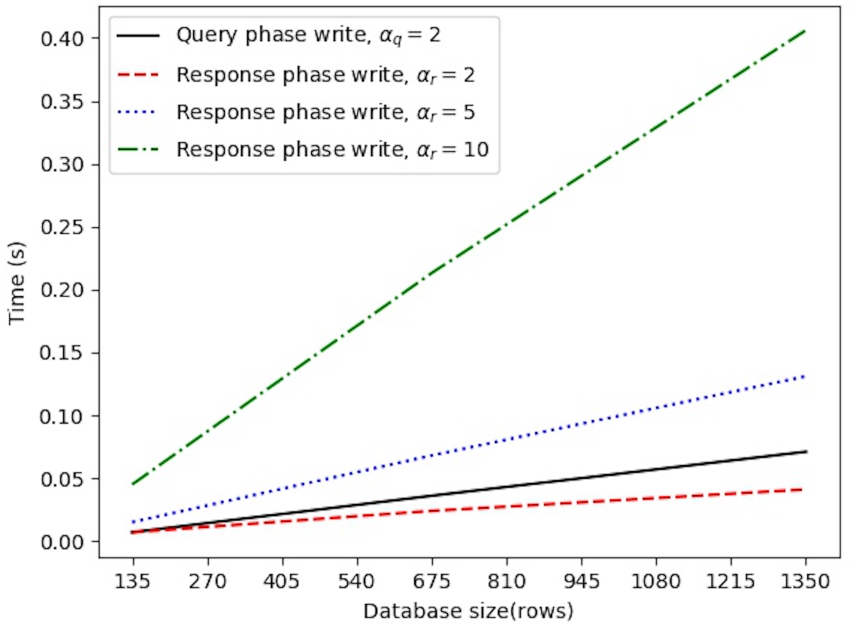
Six nodes running times for query write request, response write request with recovery from 2 collisions, response write request with recovery from 5 collisions, response write request with recovery from 10 collisions.

**Figure 6:**
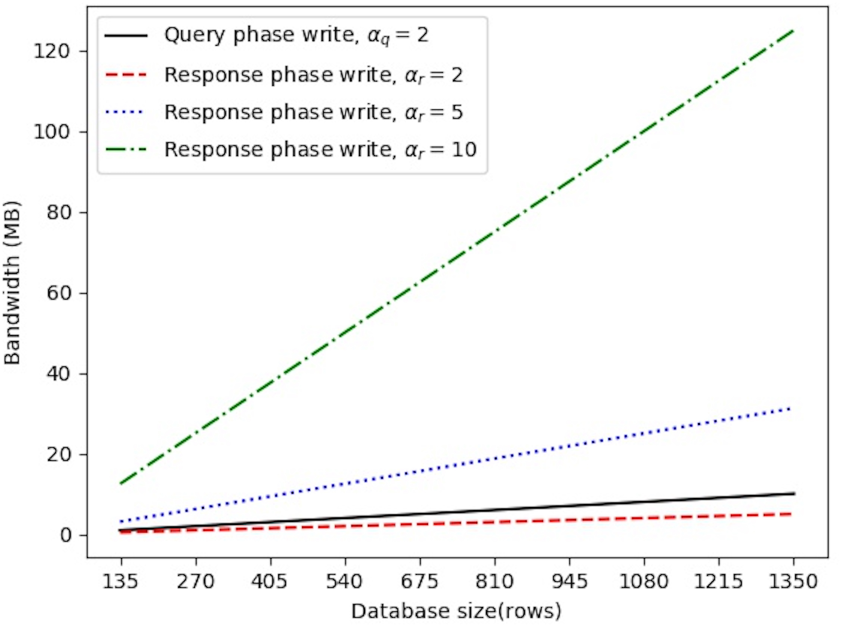
Six nodes bandwidth averages for query write request, response write request with recovery from 2 collisions, response write request with recovery from 5 collisions, response write request with recovery from 10 collisions.

On the other hand, the query phase is less efficient than the response phase (with *α*_*r*_ = 2), compared to the two-node setting, since the querying user now has to generate two vectors for each gene so that collision recovery is possible, hence, four vectors in total; whereas, the responding user only generates two vectors.

## 4 Discussion

In this section, we discuss the experimental data, as well as provide an overview of related work.

### 4.1 Remarks

Our experimental evaluation attests to the practicality of using AnoniMME to bring anonymity to the Matchmaker Exchange. Overall, using the method proposed in Section 3.3.1 to recover write requests in case of collisions yields better running times and bandwidth complexities, even when the number of nodes increases.

Since AnoniMME is based on Riposte [3], one might want to compare the two systems; however, Riposte focuses on experimental results from the server’s perspective, while we evaluate performance from a user perspective.

Also note that the bandwidth overhead in our *n*-server construction is non-negligible, especially with a high collision recovery factor and increasing database sizes (as discussed in Section 3.3.1). A possible solution would be to use distributed point functions to reduce bandwidth complexity, similar to Riposte. However, we leave this to future work.

As the anonymity set size in AnoniMME corresponds to the number of users querying in a given epoch, one could increase it by requiring users to send empty queries to the system, following a certain probability distribution. The write requests would be formed as discussed in Section 3.2, although, instead of inputting a gene, the public key, or the contact details, the users just send an empty query. This is also used in Riposte, to minimize statistical disclosure attacks on their platform.

Finally, note that our implementation currently allows for 64 character messages, thus, queries can also include phenotypes from the Human Phenotype Ontology (as currently supported by MME), although, to ease of presentation we have discussed our experiments by only considering gene names. In future work, we plan to conduct a user study simulating a real-world deployment of AnoniMME with users of the MME, aiming to evaluate its usability with respect to anonymity protection, delays introduced by epochs, etc..

### 4.2 Related Work

Rapid and effective progress in genomics and personalized medicine is often promoted as being dependent on the ability to share sequenced genomes, and make them accessible to researchers for different research purposes. However, it is often hard to share data due to privacy, ethical, legal, and informed consent hurdles. To address these issues, a few privacy-preserving methods have been presented to facilitate genomic data sharing. [7] use secret sharing for distributing data among several entities. Using secure multi-party computations on the data, computations can be done across multiple independent entities, without violating the privacy of individual donors or leaking the data to third parties. Then, [12] allow clinicians to find similar patients in bio-repositories, with similarity being defined as the edit distance. Their construction is based on a combination of a novel genomic edit distance approximation algorithm and new construction of private set difference size protocols. [2] introduce a framework using Intel’s Software Guard Extension and hardware for trustworthy computations. This way, secure and distributed computation over encrypted data is performed, respecting institutional policies and regulations for protected health information.

Another initiative developed by GA4GH, besides MME, is the Beacon Project [4]; a beacon is a service that any institution can implement to share genetic data. Users can query the system through a federated search engine, the Beacon Network. The queries are of the form *“Do you have any genomes with an ‘A’ at position 100,735 on chromosome 3?”,* and the beacon responds with either “Yes” or “No”, keeping all other sequence data concealed. This kind of queries can be used to either search all beacons or specific databases. The result is then shown as a list of databases where the allele has been previously observed, including the institution that holds that database. [10] present an attack on beacons, showing that reidentification is possible using a likelihood-ratio test. Mitigations for this attack are presented by [9], however these mitigations comes with a diminished utility of the beacon. The original attack has been improved by [11] in terms of number of queries needed to determine the presence of an individual in a beacon. Note that these attacks do not apply to MME, since no genotype information or aggregate data is released publicly, and the querying is done only on specific genes, with no genotype information.

Overall, a number of attacks to anonymized/de-identified genomic data have been presented. [6] show how to detect the presence of an individual genotype in a mixture of pooled DNA, while [5] recover the surnames of individuals from a genomic data repository by profiling short tandem repeats on the Y chromosome, querying recreational genealogy databases, and relying on metadata like age and state to recover the identity of the target.

As already mentioned, our construction is similar in nature to Riposte [3], an anonymous broadcast messaging system, which also built using Reverse PIR. Riposte allows a large number of clients to post messages anonymously on a “bulletin board” maintained at a small set of servers. The main goal of the system is to provide a platform for whistleblowers, allowing them to anonymously post 160 byte length messages. Besides using Reverse PIR in a different setting, and thus addressing different challenges in scalability, also note that our AnoniMME framework also allows replies to messages.

## 5 Conclusion

This paper presented AnoniMME, a framework geared to bring anonymity to the Matchmaker Exchange (MME) platform. Specifically, AnoniMME supports anonymous queries, by relying on Reverse PIR, while mirroring the functionalities of MME. Queries include the gene name as in MME, but also the querying user’s public key, and are collected during epochs whose length is based on the number of write requests. By taking advantage of the underlying MME matching protocol, these queries can be seamlessly responded to, without publicly revealing the contact details of other researchers/clinicians which generated a match, by using the public key provided to encrypt the match. Also, other users can provide their (encrypted) contact details if they so wish.

The proposed protocol is compatible with the functionalities and the requirements of MME, but adds anonymous queries with a low overhead, as we demonstrated empirically. Thus, we are confident that AnoniMME can eventually be deployed in the wild and further encouraging researchers to share genomic data, by minimizing the possibility of exposing confidential research when using Matchmaker Exchange.

As part of future work, we plan to include and experimentally evaluate an extension to malicious users in our prototype, support the execution of the response phase over multiple query epochs, further reduce bandwidth complexity, and perform a user study to evaluate its usability.

## Acknowledgements

We thank Christophe Dessimoz for valuable feedback provided, as well as insights from users of the platform. This work was supported by a Google Faculty Award on Enabling Progress in Genomic Research via Privacy-Preserving Data Sharing.

## References

[1] O. J. Buske et al. The Matchmaker Exchange API: Automating patient matching through the exchange of structured phenotypic and genotypic profiles. Human mutation, 36: 922–927, 2015.

[2] F. Chen et al. PRINCESS: Privacy-protecting Rare disease International Network Collaboration via Encryption through Software guard extensionS. Bioinformatics, 33: 871–878, 2017.

[3] H. Corrigan-Gibbs et al. Riposte: An Anonymous Messaging System Handling Millions of Users. In Proceedings of the 2015 IEEE Symposium on Security and Privacy, pages 321–338, 2015.

[4] Global Alliance for Genomics and Health. A federated ecosystem for sharing genomic, clinical data. Science, 352: 1278–1280, 2016.

[5] M. Gymrek et al. Identifying personal genomes by surname inference. Science (New York, N.Y.), pages 321–324, 2013.

[6] N. Homer et al. Resolving Individuals Contributing Trace Amounts of DNA to Highly Complex Mixtures Using High-Density SNP Genotyping Microarrays. PLOS Genetics, page e1000167, 2008.

[7] L. Kamm et al. A new way to protect privacy in large-scale genome-wide association studies. Bioinformatics, 29: 886–893, 2013.

[8] A. A. Philippakis et al. The Matchmaker Exchange: A platform for rare disease gene discovery. Human mutation, 36: 915–921, 2015.

[9] J. L. Raisaro et al. Addressing Beacon re-identification attacks: quantification and mitigation of privacy risks. Journal of the American Medical Informatics Association, 24: 799–805, 2017.

[10] S. S. Shringarpure and C. D. Bustamante. Privacy Risks from Genomic Data-Sharing Beacons. The American Journal of Human Genetics, 97: 631–646, 2015.

[11] N. v. Thenen et al. Re-Identification of Individuals in Genomic Data-Sharing Beacons via Allele Inference. bioRxiv, 2017.

[12] X. S. Wang et al. Efficient Genome-Wide, Privacy-Preserving Similar Patient Query Based on Private Edit Distance. In Proceedings of the 22nd ACM SIGSAC Conference on Computer and Communications Security, pages 492–503, 2015.

